# Venom Isosolenopsin A Delivers Rapid Knockdown of Fire Ant Competitors

**DOI:** 10.1101/454637

**Authors:** Eduardo G P Fox, Xiaoqing Wu, Lei Wang, Li Chen, Yong-Yue Lu, Yijuan Xu

## Abstract

Fire ant venoms are composed of insecticidal alkaloids named ‘solenopsins’. Whilst species-specific differences are reported, little attention was given to caste-specific venom adaptations. The venom of fire ants queens has remained poorly studied. Founding queens must succeed in isolation in the field, where venom is bound to play a role against competitor species. The venoms of fire ant queens are strikingly similar across different species, in being mainly composed of the alkaloid isosolenopsin A, regardless of the chemical diversity of the worker caste. From assuming this pattern as the evolutionary result of stabilising trait selection, we hypothesise a shared mechanism explaining the conserved venom composition among the fire ant queens of different species. Here we report that fire ant queen venom and its major compounds are much quicker to neutralise competitor ants than the more diverse venoms of workers. Three representative competitor ant species sympatric with invasive fire ants were selected, exposed on the head to venoms from invasive fire ant workers and queens of two main invasive species, *Solenopsis invicta* and *S. geminata*. The venom diversity in the worker caste of these species represent extremes in the chemical diversity of fire ants. Queen venoms delivers quicker knockdown of rival foragers than worker venoms. The effects are traced back to synthetic solenopsins demonstrating solenopsin A analogues are particularly efficient as contact neurotoxins. The observed effects are comparable to nicotine. Overall the venoms of *S. invicta* seem more lethal than of *S. geminata*, regardless of knockdown speed. We believe these are fundamental aspects in the chemical ecology of the invasive ants which have been long overlooked, and emphasise on the need for further studies into the venom biology of founding queens.

## 1. Introduction

Venom chemistry is shaped by the need for survival. The ecological diversity of ants is mirrored by untapped diversity of venom toxins (Touchard et al., 2016). Although ants figure amongst the most abundant extant venomous arthropods, with ca. 15,000 extant species (Antwiki, 2015), their venom chemistry is known for few species. For instance the Red Imported Fire Ant *Solenopsis invicta* Buren (henceforth RIFA) is a particularly well-studied species of remarkable aggressiveness and capable of delivering painful stings (Tschinkel, 2006). Their venoms are composed of a complex mixture of alkaloids (>90% v/v) (Fox, 2014). These alkaloids are known as ‘solenopsins’ and present marked insecticidal action (Blum, 1988; Lai et al., 2010).

As eusocial animals caste specialisation is central to ant biology. However, venom caste specialisation is a largely unexplored aspect. Fire ant workers present marked species-specific ratios of solenopsin analogues (Brand et al., 1973). However the venom of fire ant gynes (i.e. winged females, or queens) is similar across different species (Brand et al., 1973a,b; Tschinkel, 2006). Known fire ant gynes venoms capitalise on analogues of solenopin A, mainly isosolenopsin A (henceforth ‘C11’) (Brand et al., 1973a,b; Tschinkel, 2006). The reasons behind this fixed chemical configuration have remained unclear (Brand et al., 1973; Shi et al., 2015).

This investigation aimed at defining the underlying mechanisms behind the toxins composition and effects of fire ant queen venoms. We hypothesised that a stabilising trait selection impeded diversification of solenopsins composition in the queen caste across the different fire ant species. In such a scenario queen venoms would be playing a central role in fire ant biology.

It is known that nest founding is the most fragile stage in the life cycle of fire ant colonies (Tschinkel, 2006). This is when newly mated queens coming from a nuptial flight must seek shelter to found a new colony. They must survive for ca. 5 weeks (Tschinkel, 2006; Markin et al., 1972) in the absence of workers to protect them. During this period rival ant species are a major obstacle for the establishment of new fire ant colonies (Vinson & Rao, 2004; Rao & Vinson, 2004). Our investigations demonstrate: (i) a remarkable knockdown effect from intoxication by fire ant gynes against rival ant species, (ii) that the major compound isololenopsin A is responsible for most of the observed effects.

## 2. Materials and Methods

### 2.1. Animals

Colonies of the crazy ant *Paratrechina longicornis* L., the acrobat ant *Crematogaster dohrni* Mayr, and the ghost ant *Tapinoma* nr. *melanocephalum* F. were obtained from a fire ant-invaded area in Guangdong, P.R. China. These species are locally abundant, and represent ant genera typically observed associated in fire ant-infested areas elsewhere, e.g. in South America (personal experience of EGPF). Mature colonies of RIFA and *Solenopsis geminata* F. (TIFA) were obtained from Beihai, Guangxi, P.R. China and separated ifrom soil according with methods described in details elsewhere (e.g. Fox et al. 2013).

These ant colonies were conditioned inside ca. 50 × 25 cm plastic boxes with internal walls painted with anti adherent Fluon. They were given adult commercial *Locusta migratoria* L. every two or three days and provided with water ad libitum. Colonies were reared at fluctuating room conditions between 25-30°C and relative humidity 55-90%, photoperiod L:D 14:10.

### 2.2. Venom extraction, purification, fractions

Venom from fire ants was extracted and fractions from *S. invicta* workers and gynes obtained following methods detailed elsewhere (Fox et al. 2013, Shi et al. 2015, respectively). In short, colonies maintained in the laboratory as described above were first inspected to manually sort numerous (>1,000) gynes and workers (>60,000). The sorted gynes and workers were immersed in pure HPLC-grade *n*-hexane with 1:5 water for 12h (after Fox et al. 2013), after which the solvent was recovered and concentrated for later purification of alkaloids and fractions through graded elutions with hexane : acetone in silica-gel columns (Shi et al. 2015). Through this method we obtained about 600 mg of pure alkaloids mixture, and about 200 mg of purified cis and trans fractions of each caste.

### 2.3. GC-MS analysis

Crude venom samples of tested fire ants were sampled as a single droplet applied to the inner wall of a 0.5 mm ID capillary tube, dissolved into 10 μg/mL nicotine in hexane as internal standard solution for relative dosage of peaks. Likewise a single droplet of each of venom fractions (purified alkaloids, *cis*- and *trans*-alkaloids fraction) was collected for relative analysis of peaks. GC-MS injections were performed as described in Fox et al. (2017) into an Agilent GC-7890B system coupled with an MS-5977B MSD by manual 1-μl injections. Obtained chromatograms were analysed with OpenChrom 1.2.0 Community Edition. External controls were nicotine and a mixture of hydrocarbons. Obtained GC-MS chromatograms and the method files can be found among Supplementary Files (Fox et al. 2018).

### 2.4. Synthetic alkaloids

Synthetic racemic solenopsins (Table S1) were purchased from WuXi AppTec (ShangHai, China) after synthesis according with Pianaro et al (2012) for cis isomers and Herath & Nanayakkara (2008) for trans isomers. Synthetic racemic nicotine was purchased from Sigma. ‘Synthetic venoms’ were produced mimicking natural venoms by mixing the main components (those present at >10% of the natural extract) present in gynes’ venoms using precision syringes according with their relative proportions observed in *S. invicta* and *S. geminata*.

### 2.5. Venom bioassays

#### 2.5.1. Venom application

Methods were designed to simulate effects from topical applications of venom. It should be emphasized that fire ants cannot introduce the stinger into the body of other hard-bodied insects of similar size because of insufficient leverage as they must bite to sting. Therefore fire ants must fight other ants using mandibles and their venom, usually by contact application.

Ants in this study were anesthetized prior to manipulation using either CO_2_ (*P. longicornis, T. melanocephalum*) or placing on ice packs (*C. rogenhoferi, Solenopsis* spp.). Dedicated description of the manipulation of each species is detailed in the Supplementary Files (Fox et al. 2018).

The effect of venoms of fire ant workers and gynes on other ants were assessed using two different approaches: (i) direct single application from touching with live excised gasters, and (ii) using venom alkaloid fractions obtained as described in Shi et al. (2015). Droplets of venom fractions and synthetic compounds were applied using a finely stretched capillary tube attached with silicon to a glass precision 1-μl syringe (See Supplementary Files videos and images for details, Fox et al. 2018). Single droplets were used calibrated under a stereomicroscope with a micrometric eyepiece to ca. 20-30 nl as estimated from diameter while gently pressing the syringe plunger. Volume set is based on Brand et al. (1974a) and measured pictures of venom droplets. Droplets were applied either on the head, typically between the frons and mouthparts, unless stated otherwise; e.g. antennae. Intoxication was assessed by perceived altered behaviour (e.g. rubbing, self-venom application, agitation) which was always followed by a state of shaking paralysis, knockdown (section 2.6), and ultimately death (section 2.6).

### 2.6. Tests for knockdown times (KDt)

Assayed ants were placed individually in vertically-supported plastic eppendorf tubes (n=10) containing a 0.8 cm-wide rectangular strip of paper provided as a standing substrate. Two digital timers were set from the onset until the end of the tests from which individual KDt times laps were noted every time an assayed ant completely lost contact with substrate, falling down to the bottom of the tube. At the end of 60 mins all assayed ants were transferred to in a Petri dish as described below to monitor for mortality after 12 h.

### 2.7. Tests for mortality

Ants of either the crazy ant, ghost ant, or acrobat ants to be assayed were grouped (n = 10) inside 7-cm wide acrilate Petri dishes provided with 2 mL drinking water pressed with cotton into an centrifuge tube, with a 20 cm^2^ piece of filter paper a standing and sheltering substrate. Their reactions were observed for 1h under a ZEISS stereomicroscope followed by a rapid check every 2 h until 12 h. Plates were tipped to the side upon every check and any paralysed individuals failing to hang onto the substrate paper were recorded as either “knocked down" (when still moving) or “dead” (when clearly not moving). Negative control ants included ants merely touched with a glass needle containing with no chemicals and ants touched with a droplet of acetone. Positive controls included ants tested with synthetic nicotine and synthetic solenopsins.

### 2.8. Imaging

Pictures of intoxicated ants, venom droplets were taken with a stereomicroscopes ZEISS Stereo Discovery v. 20 for images. Videos were taken with a Miniso macro lens clipped to an iPhone 6. Venom droplet sizes were measured with MeasurePictures v.1.0 and volume estimated by extrapolation to a perfect sphere using the formula

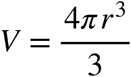

Videos were edited with iMovie 10 using a MacBook Pro Retina mid-2015. Obtained images can be found in the Supplementary Files.

### 2.9. Statistical analysis

All numeric data was input into R v. 3.2.2 for graphical representation of plotted results and for general statistical analysis. The following R packages were used: ‘plyr’, ‘reshape2’, ‘ggplot2’, ‘HMRd’. Numbers of dead individuals were compared by non-parametric Kruskal-Wallis at alpha = 0.05 complemented with Dunn’s paired test. Scripts containing all the plots details and raw data are provided in Supplementary Files (Fox et al. 2018).

## 3. Results

Competitor ant species showed intoxication symptoms shortly after contact with fire ant venom, represented in Table 1. Observed initial effects generally confirm observations by previous authors from competitors intoxication with fire ant worker venoms (Greenberg et al., 2008; LeBrun et al., 2014).

**Table 1.** Perceived behavioural alterations within 1h following topical application of venom, venom fractions and synthetic compounds from fire ant workers and gynes on three different species of competitor ants, translated as tentative frequencies from annotations taken during the observations. Short videos exemplifying of these behaviours are provided as Supplementary Materials (Fox et al. 2018).

|  | Agitation | Rubbing | Venom dabbing | Grooming | No difference |
| --- | --- | --- | --- | --- | --- |
| **Crude Venom** |  |  |  |  |  |
| Gyne *S. geminata* | + | + |  | + | +++ |
| Gyne *S. invicta* | + | ++ | + | ++ |  |
| Worker *S. geminata* | + | + | + | ++ | + |
| Worker *S. invicta* |  | ++ | ++ | ++ |  |
| **Cis Fraction** |  |  |  |  |  |
| Extract gynes *S. invicta* | + | ++ | + | ++ |  |
| Extract workers *S. invicta* | + |  |  | ++ | +++ |
| C11 |  | ++ | ++ | ++ |  |
| C13 | +++ |  | + |  | ++ |
| C15 | +++ |  |  |  | +++ |
| **Trans fraction** |  |  |  |  |  |
| Extract gynes *S. invicta* |  | ++ |  | ++++ |  |
| Extract workers *S. invicta* | + | +++ | + | + |  |
| C11 | + | +++ | ++ |  |  |
| C13 |  | +++ | ++ | + |  |
| C15 | + | ++ |  | + | ++ |
| Nicotine | + | ++ | ++ | + |  |

Notes: *Paratrechina longiconis* seemed to actively avoid application of gynes venom and cis-C11; *P. longicornis* and *S. invicta* workers would often stay motionless for few seconds after application of *S. invicta* gynes’ venom and *S. geminata* workers’ venom; *Crematogaster* cf. *rogenhoferi* workers would often display spontaneous exudation of fluids (i.e. venom and vomiting leaving yellow stains on filter paper) when paralyzed after 1h from contact with *S. invicta* workers venom, C13 isomers and trans C15.

Figure 1 however demonstrates envenomation by fire ant queens venoms usually leads to rapid knockdown (also see Supplementary videos in Fox et al. (2018)), which takes longer or may not happen following worker venom exposure (Fig. 1A). A higher speed of knockdown is not directly related with increased mortality later (Fig. 1B).

**Fig. 1.**
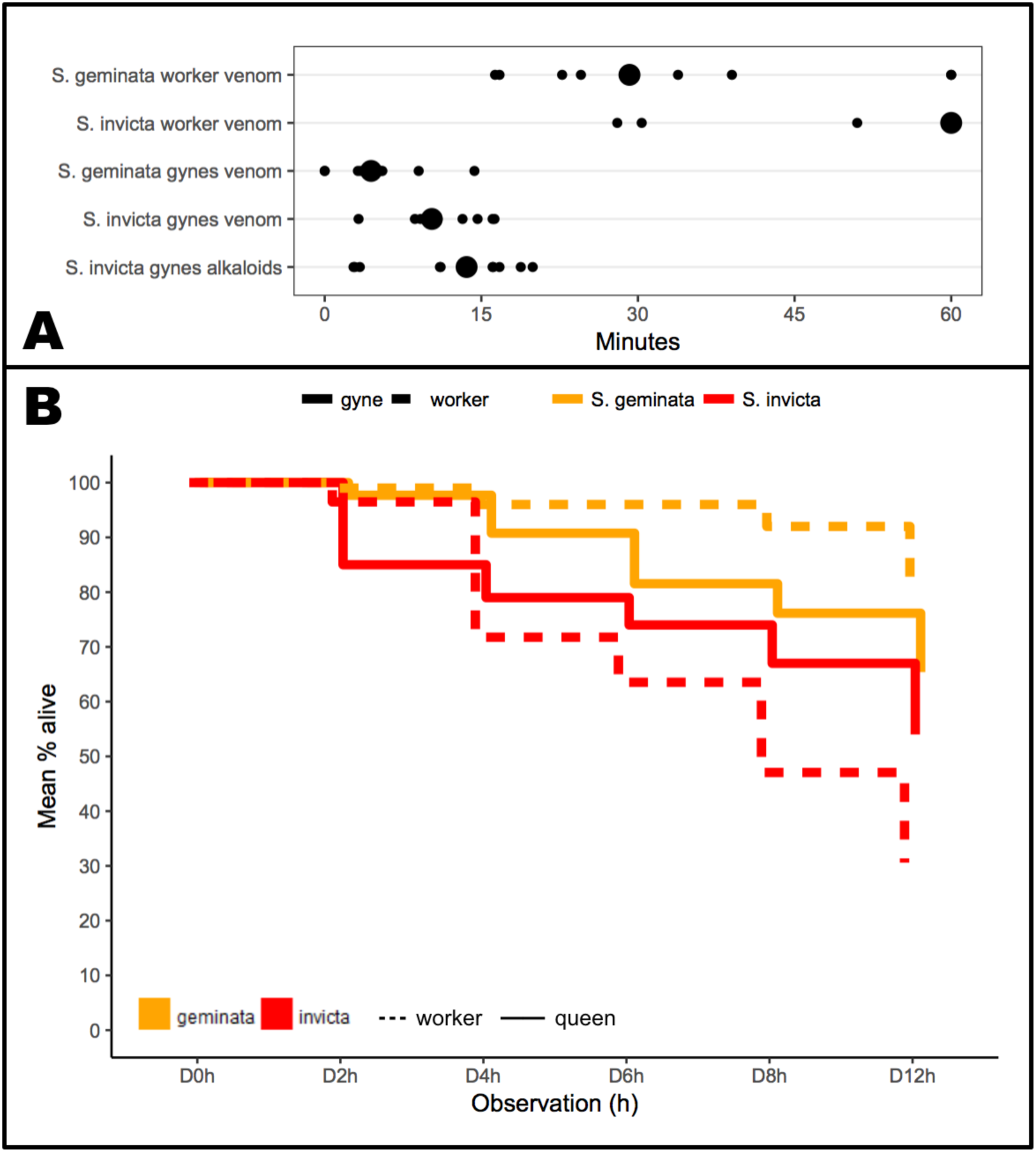
Speed of knockdown (A) and mean mortality (B) among workers of the crazy ant *Paratrechina longicornis* exposed on the head contact to a droplet of invasive fire ant venoms in natura. Statistical analysis: (A) Quickest knockdown is delivered by queens of *Solenopsis geminata* (W = 86, p-value = 0.007); slowest knockdown is delivered by *S. invicta* workers (W = 0, p-value < 0.001); *S. invicta* crude venom has same effect as pure alkaloids. (B) The venom of *S. geminata* was over time the least lethal of all (p-values = 0.001; 0; 0.001) based on Kruskal-Wallis test, chi-squared = 14.254, df = 3.; the venom of *S. invicta* workers was the most lethal after 12 h (p-values = 0, 0.001, 0), chi-squared = 22.438, df = 3. Mortality from gynes’ venoms was overall equivalent.

Both species produce the same diversity range of venom solenopsins (Fig.2) however the relative proportion of trans solenopsins isomers, particularly *trans*-C13 (solenopsin B) and *trans*-C15 (solenopsin C) are greater in RIFA *S. invicta*. The venoms of queens capitalise on *cis*- and *trans*-C11 (Fig. 2). These observations are in accordance with the reported by previous authors (e.g. Shi et al. 2014).

**Fig. 2.**
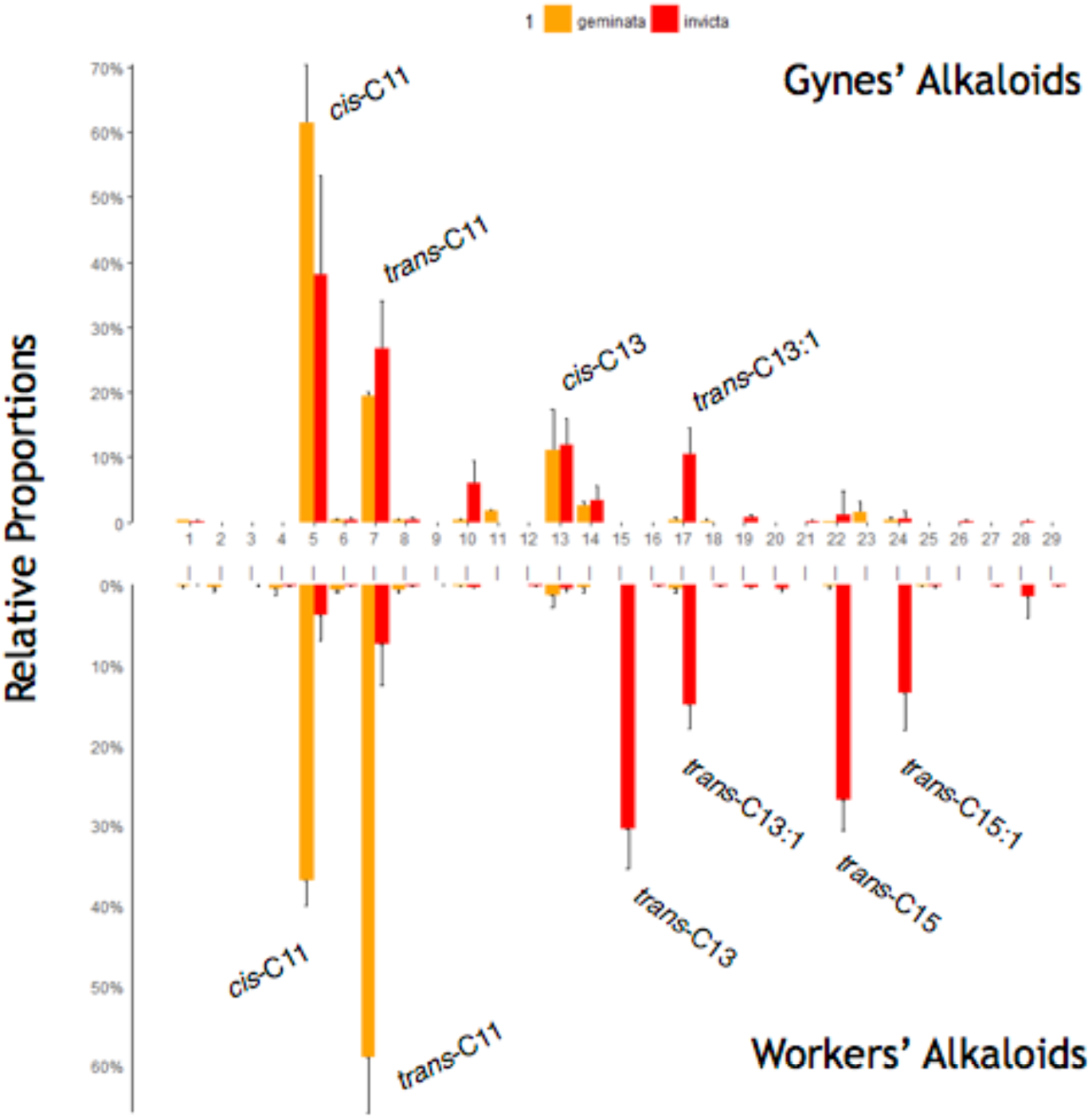
Relative proportions (as % means ± standard deviations) of venom alkaloids across queens (gynes) and workers in the fire ants *Solenopsis invicta* and *S. geminata* as identified by GC-MS analysis of venom droplets from individual insects (n = 5). Numbers correspond to compounds listed in Table S1. Saturated compounds were synthesised to compare for insecticidal effects; unsaturated solenopsin analogues are more difficult to obtain, and shall be dealt with in a subsequent study focusing on the venom of workers.

Synthetic solenopsins were used to reproduce the observed intoxication effects. Knockdown times (KDt) by synthetic isosolenopsin A (*cis*-C11) were significantly shorter than other compounds. Isomers of solenopsin C (C15) mostly did not cause a knockdown (Fig. 3) for most tests, however an absence of knockdown within 1h does not imply intoxicated ants had recovered.

**Fig. 3.**
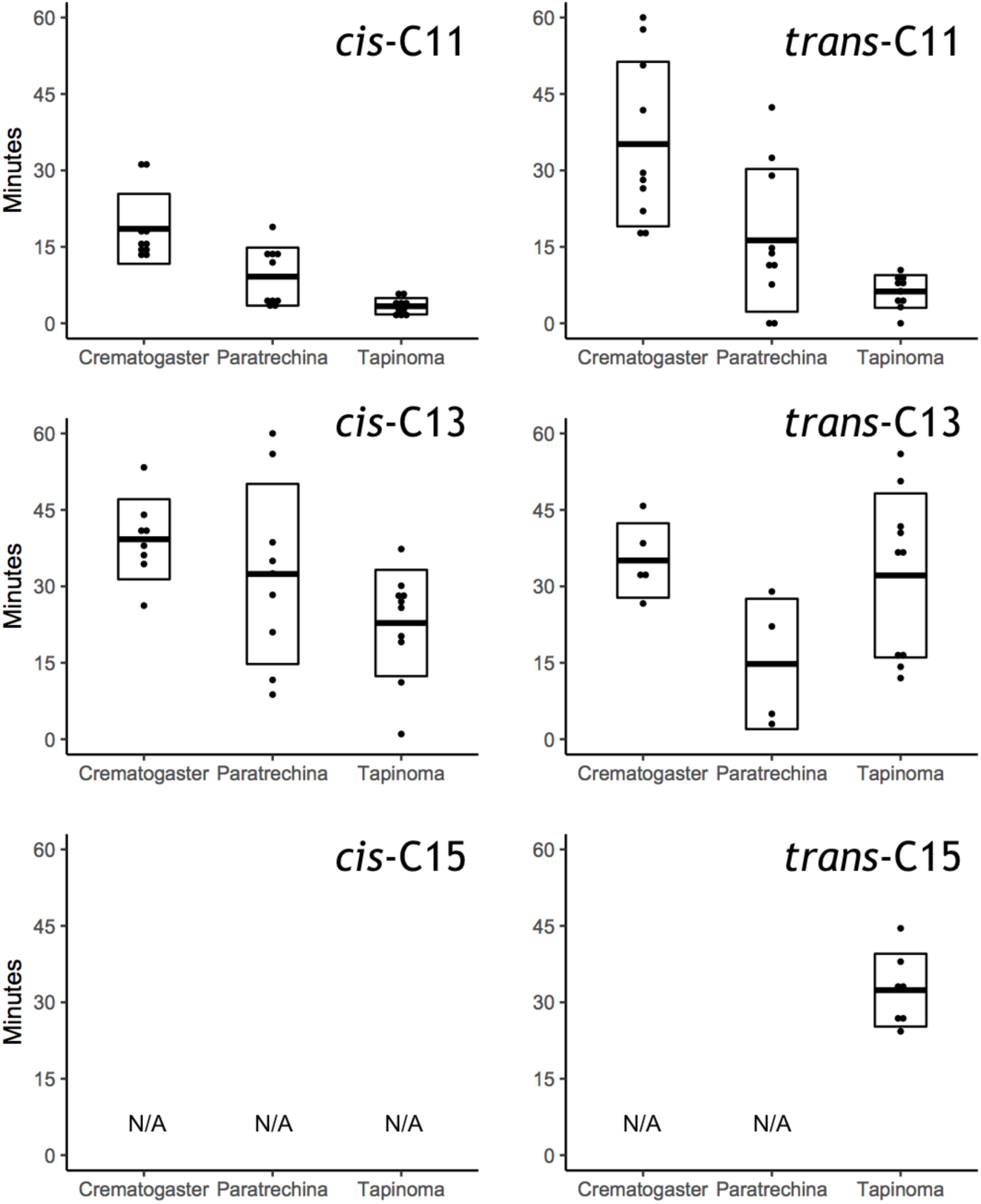
Knockdown speed from the application of a 20-30 µl droplet of synthetic solenopsins to the heads of ant workers of three representative competitor species. Box plots represent mean ± SD among groups of 10 treated workers monitored for 1 h. Statistical Analysis: Isomers of C11 had significantly different effect on the tested species (Kruskal-Wallis chi-squared = 19.4416, df = 2, p-value = 0); while C13 had similar overall effect regardless of isomer or species. Measured knockdown speed delivered by *ci*s-C11 was the shortest of all solenopsins regardless of species (Kruskal-Wallis chi-squared = 19.4416, df = 2, p-value = 0).

Mixtures of C11 analogues and C13 were concocted to mimic the venom composition of TIFA and RIFA queens venoms, and largely reproduced observed effects (Fig. 4). Comparison with natural alkaloid fractions further confirmed the observed effects are only due to venom solenopsins (Fig. S1). Rapid knockdown from the venom of fire ant gynes therefore chiefly derives from intoxication by isosolenopsin A, *cis*-C11. Although mortality following contact with pure solenopsins is stronger, the general pattern is maintained (compare Fig. 1 and Fig. 4). Other solenopsin analogues may be associated with higher mortality, particularly of isosolenopsin B (*cis*-C13) (see Fig. S2), possibly accounting for higher mortality in RIFA venoms (Fig. 1). The observed effects were visually similar (albeit slower) to intoxication by the insecticidal alkaloid nicotine (see Supplementary video), suggesting that intoxication by these alkaloids follows related pathways.

**Fig. 4.**
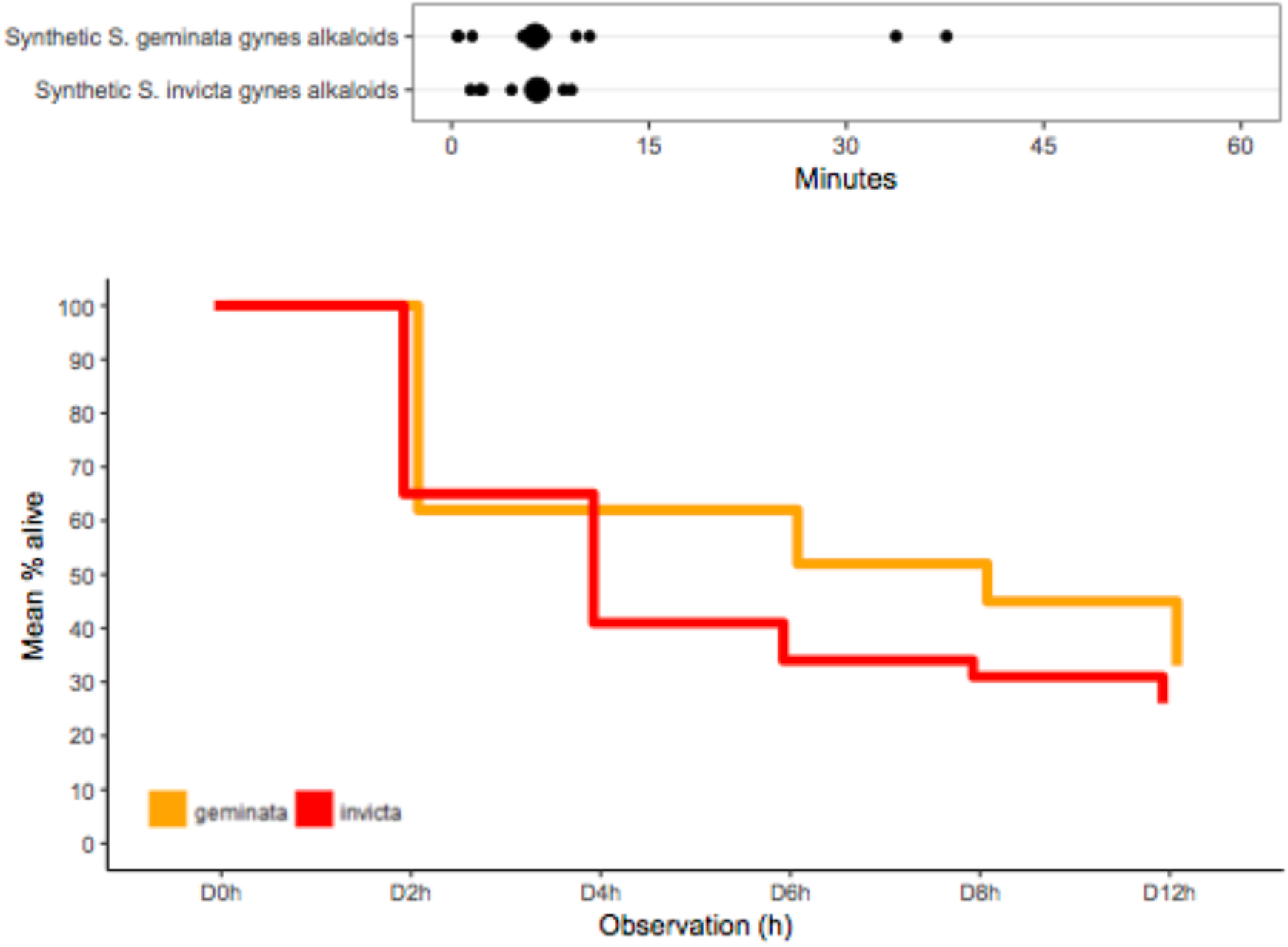
Speed of knockdown (A) and mean mortality (B) among workers of the crazy ant *Paratrechina longicornis* exposed on the head contact to a 20-30 µl droplet of a mixture of synthetic solenopsins mimicking the venoms of invasive fire ant queens. Statistical Analysis: (A) & (B) Overall the effects of both mixtures of solenopsins were similar. (B) The mortality at 4h was greater for the venom of *S. invicta*.

Because fire ants are larger than workers and have a prominently swollen gaster, they are physically unable to gaster-flag or inject venom like workers do. Even when encouraged to sting into prey or skin a gyne’s stinger will not penetrate (personal observations by EGPF). Therefore gynes are apparently limited to using venom via topical application, either on eggs during oviposition (Tschinkel, 2006) or by attacking other arthropods. Our results demonstrate gynes can quickly neutralise competitor ant foragers. This trait is likely invaluable during claustral nest foundation, which is a lifecycle challenge shared by all fire ant species, as is the composition of their venoms. We are currently working on simulating field conditions under controlled conditions, and verifying the effects of solenopsins as repellents an antimicrobials as possible additional defence against natural enemies. In conclusion the present investigation demonstrates for the first time that the venom of fire ant gynes is particularly effective for the rapid neutralisation of competitor ants, apparently tuned within strict proportions of analogous alkaloids. We believe this is an overlooked aspect in the biology of fire ants which helps understand the invasive success of this clade.

## Acknowledgements

Wen-Wei Tang, HuiJu Zhang, Meng Xu, Mu-Yang He, Dong-Qiang-Zeng provided invaluable support in collecting fire ant colonies in the field. Kevin Haight provided useful insights into venom manipulation. Useful comments were provided by Laurent Keller and Edward LeBrun. The present and last publication complied with unique local requirements of South China Agricultural University, Guangdong, China regarding certain ownerships, to be discussed elsewhere.

## References

Antwiki (2015) http://www.antwiki.org/wiki/Welcome_to_AntWiki

Blum, M.S. (1988) Biocidal and deterrent activities of nitrogen heterocycles produced by venomous myrmicine ants, pp. 438‹449. In H. G. Cutler [ed.], Biologically active natural products: potential use in agriculture. American Chemical Society Symposium Ser. 380, Washington, DC.

Brand, J.M., Blum, M.S., and Barlin, M.R. (1973a) Fire ant venoms: intraspecific and interspecific variation among castes and individuals. Toxicon 11,325–331.

Brand, J.M., Blum, M.S., and Ross, H.H. (1973b) Biochemical evolution in fire ant venoms. Insect Biochem. 3,45–51.

Fox E.G.P. Venom toxins of fire ants. In: Gopalakrishnakone P., Calvete J.J., editors. Venom genomics and proteomics. Springer; Dordrecht, The Netherlands: 2014. pp. 1–16.

Fox E.G.P., Pianaro A, DR Solis, Delabie JHC, Vairo BC, Machado EA, Correa OC. (2012) Intraspecific and Intracolonial Variation in the Profile of Venom Alkaloids and Cuticular Hydrocarbons of the Fire Ant *Solenopsis saevissima* Smith (Hymenoptera: Formicidae). Psyche; 2012: 1–10.

Fox, E.G.P., Solis, D.R., Santos, L.D., Pinto, J.R.A.S., Menegasso, A.R.S., Silva, R.,C.M.C., Palma, M.S., Bueno, O.C., Machado, E.A. (2013). A simple, rapid method for the extraction of whole fire ant venom (Insecta: Formicidae: *Solenopsis*). Toxicon 65, 5–8. https://doi.org/10.1016/j.toxicon.2012.12.009

Fox, E.G.P., Chen, L. XiaoQing, W. (2018) On the Venom Solenopins alkaloids of Fire Ants: Beware Lonely Queens. http://dx.doi.org/10.17632/s8x82r464b.1

Greenberg, L., Kabashima, J.N., Allison, C.J., et al (2008) Lethality of Red Imported Fire Ant Venom to Argentine Ants and Other Ant Species. Ann Entomol Soc Am 101:1162–1168. doi: 10.1603/0013-8746-101.6.1162

Herath, H.M.T.B., Nanayakkara, N.P.D. (2008) Synthesis of enantiomerically pure fire ant venom alkaloids: Solenopsins and lsosolenopsins A, B and C. J. Heterocyc. Chem. 45: 129–136. https://doi.org/10.1002/jhet.5570450112

Lai L.C.Y.Y., Chang R.N., Huang C.U., Wu J.W. (2010) Comparative toxicity of two fire ant venoms to *Spodoptera litura* larvae. Sociobiology; 56: 653–663.

LeBrun, E.G.; Jones N.T.; Gilber L.E. (2014) “Chemical Warfare Among Invaders: A Detoxification Interaction Facilitates an Ant Invasion”. Science. 343 (6174): 1014–1017.

Markin G.P., Collins H.L., Dillier J.H. (1972) Colony founding by queens of the red imported fire ant, *Solenopsis invicta*. Ann Entomol Soc Am 65:1053–1058

Pianaro, A., Fox, E.G.P., Bueno, O.C., Marsaioli, A.J. (2012) Rapid configuration analysis of the solenopsins. Tetrahedron Asymmetry 23, 635–642. https://doi.org/10.1016/j.tetasy.2012.05.005

Rao A., Vinson S.B. (2004) Ability of resident ants to destruct small colonies of *Solenopsis invicta* (Hymenoptera : Formicidae). Environ Entomol 33:587–598. doi: http://www.entsoc.org/pubs/periodicals/ee/index.htm

Shi, Q.H., Hu, L., Wang, W.K., Vander Meer, R.K., Porter, S.D., Chen, L. (2015) Workers and alate queens of *Solenopsis geminata* share qualitatively similar but quantitatively different venom alkaloid chemistry. Front. Ecol. Evol. 3, 76

Touchard A., Aili S.R., Fox E.G.P., Escoubas P., Orivel J., Nicholson G.M., Dejean A. (2016) The Biochemical Toxin Arsenal from Ant Venoms. Toxins, 20: 8 (1) pii: E30. doi: 10.3390/toxins8010030.

Tschinkel W.S. The Fire Ants. (2006) Cambridge: Belknap Press of Harvard University Press.

Vinson S.B., Rao A. (2004) Inability of incipient *Solenopsis invicta* (Hymenoptera: Formicidae) colonies to establish in a plot with a high density of Solenopsis (*Diplorhoptrum*) colonies. Environ Entomol 33:1626–1631. DOI: 10.1603/0046-225X-33.6.1626

